# Genotypic variability enhances the reproducibility of an ecological study

**DOI:** 10.1101/080119

**Authors:** Alexandru Milcu, Ruben Puga-Freitas, Aaron M. Ellison, Manuel Blouin, Stefan Scheu, Thomas Girin, Grégoire T. Freschet, Laura Rose, Michael Scherer-Lorenzen, Sebastien Barot, Jean-Christophe Lata, Simone Cesarz, Nico Eisenhauer, Agnès Gigon, Alexandra Weigelt, Amandine Hansart, Anna Greiner, Anne Pando, Arthur Gessler, Carlo Grignani, Davide Assandri, Gerd Gleixner, Jean-François Le Galliard, Katherine Urban-Mead, Laura Zavattaro, Marina E.H. Müller, Markus Lange, Martin Lukac, Michael Bonkowski, Neringa Mannerheim, Nina Buchmann, Olaf Butenschoen, Paula Rotter, Rahme Seyhun, Sebastien Devidal, Zachary Kayler, Jacques Roy

**Author notes:** Corresponding author: Alexandru Milcu, CNRS, Ecotron - UPS 3248, Campus Baillarguet, 34980, Montferrier-sur-Lez, France.

## Abstract

Many scientific disciplines currently are experiencing a “reproducibility crisis” because numerous scientific findings cannot be repeated consistently. A novel but controversial hypothesis postulates that stringent levels of environmental and biotic standardization in experimental studies reduces reproducibility by amplifying impacts of lab-specific environmental factors not accounted for in study designs. A corollary to this hypothesis is that the deliberate introduction of controlled systematic variability (CSV) in experimental designs can increase reproducibility. We tested this hypothesis using a multi-laboratory microcosm study in which the same ecological experiment was repeated in 14 laboratories across Europe. Each laboratory introduced environmental and genotypic CSV within and among replicated microcosms established in either growth chambers (with stringent control of environmental conditions) or glasshouses (with more variable environmental conditions). The introduction of genotypic CSV led to lower among-laboratory variability in growth chambers, indicating increased reproducibility, but had no significant effect in glasshouses where reproducibility also was lower. Environmental CSV had little effect on reproducibility. Although there are multiple causes for the “reproducibility crisis”, deliberately including genetic variation may be a simple solution for increasing the reproducibility of ecological studies performed in controlled environments.

---

Reproducibility—the ability to duplicate a study and its findings—is a defining feature of scientific research. In ecology, it is often argued that it is virtually impossible to accurately duplicate any single ecological experiment or observational study. The rationale is that the complex ecological interactions between the ever-changing environment and the extraordinary diversity of biological systems exhibiting a wide range of plastic responses at different levels of biological organization make exact duplication unfeasible^1,2^. Although this may be true for observational and field studies, numerous ecological (and agronomic) studies are carried out with artificially assembled simplified ecosystems and controlled environmental conditions in experimental microcosms or mesocosms (henceforth, “microcosms”)^3–5^. Since biotic and environmental parameters can be tightly controlled in microcosms, results from such studies should be easier to reproduce. Even though microcosms have frequently been used to address fundamental ecological questions^4,6,7^, there has been no quantitative assessment of the reproducibility of any microcosm experiment.

Experimental standardization—the implementation of strictly defined and controlled properties of organisms and their environment—is widely thought to increase both reproducibility and sensitivity of statistical tests^8,9^ because it reduces within-treatment variability. This paradigm has been recently challenged by several studies on animal behavior, suggesting that stringent standardization may, counterintuitively, be responsible for generating non-reproducible results^9–11^ and contribute to the actual reproducibility crisis^12–15^; the results may be valid under given conditions (i.e., they are local “truths”) but are not generalizable^8,16^. Despite rigorous adherence to experimental protocols, laboratories inherently vary in many conditions that are not measured and are thus unaccounted for, such as experimenter, micro-scale environmental heterogeneity, physico-chemical properties of reagents and lab-ware, pre-experimental conditioning of organisms, and their genetic and epigenetic background. It even has been suggested that attempts to stringently control all sources of biological and environmental variation might inadvertently lead to the amplification of the effects of these unmeasured variations among laboratories, thus reducing reproducibility^9–11^.

Some studies have gone even further, hypothesizing that the introduction of controlled systematic variation (CSV) among the replicates of a treatment (e.g., using different genotypes or varying the organisms’ pre-experimental conditions among the experimental replicates) should lead to less variable mean response values between the laboratories that duplicate the experiments^9,11^. In short, it has been argued that reproducibility should increase by shifting the variance from among experiments to within them^9^. If true, then introducing CSV will increase researchers’ ability to draw generalizable conclusions about the directions and effect sizes of experimental treatments and reduce the probability of false positives. The trade-off to this approach is that increasing within-experiment variability will reduce the sensitivity (i.e. the probability of detecting true positives) of statistical tests. However, it currently remains unclear whether introducing CSV increases reproducibility of ecological microcosm experiments, and if so, at what cost for the sensitivity of statistical tests.

To test the hypothesis that introducing CSV enhances reproducibility in an ecological context, we had 14 European laboratories simultaneously run a simple microcosm experiment using grass (*Brachypodium distachion* L.) monocultures and grass and legume (*Medicago truncatula* Gaertn.) mixtures. As part of the reproducibility experiment, the 14 laboratories independently tested the hypothesis that the presence of the legume species *M. truncatula* in mixtures would lead to higher total microcosms plant productivity and enhanced growth of the non-legume *B. distachion via* rhizobia-mediated nitrogen fertilization and/or nitrogen sparing effects^17–19^.

All laboratories were provided with the same experimental protocol, seed stock from the same batch, and identical containers in which to establish microcosms with grass only and grass-legume mixtures. Alongside a control (CTR) with no CSV and containing a homogenized soil substrate (mixture of soil and sand) and a single genotype of each plant species, we explored the effects of five different types of within- and among-microcosm CSV on experimental reproducibility of the legume effect (Fig. 1): 1) within-microcosm environmental CSV (ENV_W_) achieved by spatially varying soil resource distribution through the introduction of six sand patches into the soil; 2) among-microcosm environmental CSV (ENV_A_), which varied the number of sand patches (none, three or six) among replicate microcosms; 3) within-microcosm genotypic CSV (GEN_W_) that used three distinct genotypes per species planted in homogenized soil in each microcosm; 4) among-microcosm genotypic CSV (GEN_A_) that varied the number of genotypes (one, two or three) planted in homogenized soil among replicate microcosms; and 5) both genotypic and environmental CSV (GEN_W_+ENV_W_) within microcosms that used six sand patches and three plant genotypes per species in each microcosm. In addition, we tested whether CSV effects depended on the level of standardization within laboratories by using two common experimental approaches (‘SETUP’ hereafter): growth chambers with tightly controlled environmental conditions and identical soil (eight laboratories) or glasshouses with more loosely controlled environmental conditions and different soils (six laboratories; see Supplementary Table 1 for the physicochemical properties of the soils).

**Fig. 1.**
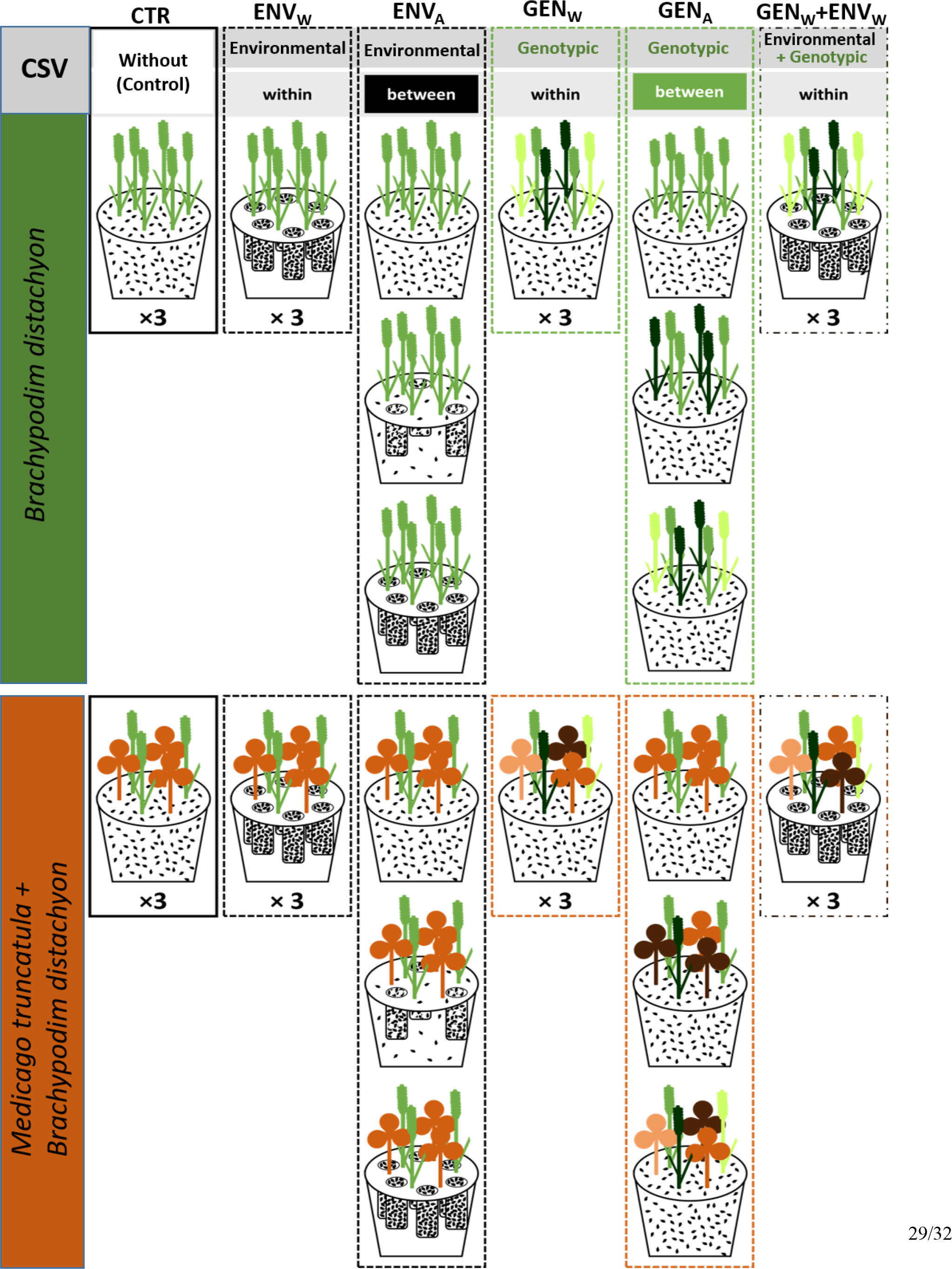
Experimental design of one block. Grass monocultures of *Brachypodium distachyon* (green shades) and grass-legume mixtures with the legume *Medicago trunculata* (orange-brown shades) were established in 14 laboratories; shades of green and orange-brown represent three distinct genotypes of *B. distachyon* (Bd21, Bd21-3 and Bd3-1) and *M. truncatula* (L000738, L000530 and L000174). Plants were established in a substrate with equal proportions of sand (black spots) and soil (white), with the sand being either mixed with the soil or concentrated in sand patches to induce environmental controlled systematic variability (CSV). Combinations of three distinct genotypes were used to establish genotypic CSV. Alongside a control (CTR) with no CSV and containing one genotype (L000738 and/or Bd21) in a homogenized substrate (soil-sand mixture), five different types of environmental or genotypic CSV were used as treatments: 1) within-microcosm environmental CSV (ENV_W_) achieved by spatially varying soil resource distribution through the introduction of six sand patches into the soil; 2) among-microcosm environmental CSV (ENV_A_), which varied the number of sand patches (none, three or six) among replicate microcosms; 3) within-microcosm genotypic CSV (GEN_W_) that used three distinct genotypes per species planted in homogenized soil in each microcosm; 4) among-microcosm genotypic CSV (GEN_A_) that varied the number of genotypes (one, two or three) planted in homogenized soil among replicate microcosms; and 5) both genotypic and environmental CSV (GEN_W_+ENV_W_) within microcosms that used six sand patches and three plant genotypes per species in each microcosm. The “× 3” indicates that the same genotypic and sand composition was repeated in three microcosms per block. The spatial arrangement of the microcosms in each block was re-randomized every two weeks. The blocks represent two distinct chambers in growth chamber setups, whereas in glasshouse setups the blocks represent two distinct growth benches in the same glasshouse.

As response variables we measured 12 parameters representing a typical ensemble of variables measured in plant-soil microcosm experiments. Six of these measured at the microcosm-level: shoot biomass, root biomass, total biomass, shoot to root ratio, evapotranspiration and decomposition of a common substrate using a simplified version of the “teabag litter decomposition method”^20^. The other six were measured on the *B. distachyon* grass species: seed biomass, height and shoot tissue chemistry including N%, C%, δ^15^N, δ^13^C. All 12 variables were then used to calculate the effect of the presence of a nitrogen-fixing legume on ecosystem functions in grass-legume mixtures (‘net legume effect’ hereafter) (Supplementary Table 2) calculated as the difference between the values measured in the microcosms with and without legumes, an approach often used in legume-grass binary cropping systems^19,21^ and biodiversity-ecosystem function experiments^17,22^.

Because we considered that statistically significant differences among the 14 laboratories would indicate a lack of reproducibility, we first assessed how our experimental treatments (CSV and SETUP) affected the number of laboratories that produced results that could be considered to have reproduced the same finding. We then determined how experimental treatments affected standard deviation (SD) of the legume effect for each of the 12 variables both within- and among-laboratories; lower among-laboratory SD implies that the results were reproduced more closely. Lastly, we explored the relationship between within- and among-laboratory SD as well as how the experimental treatments affected the statistical power of detecting the net legume effect.

Although each laboratory followed the same experimental protocol, we found a remarkably high level of among-laboratory variation for the majority of response variables (Supplementary Fig. 1) and the net legume effect on those variables (Fig. 2). For example, the net legume effect on mean total plant biomass varied among laboratories from 1.31 to 6.72 g dry weight (DW) per microcosm in growth chambers, suggesting that unmeasured laboratory-specific conditions outweighed effects of experimental standardization. Among glasshouses, differences were even larger: the legume effect on the mean plant biomass varied by two orders of magnitude, from 0.14 to 14.57g DW per microcosm (Fig. 2). Furthermore, for half of variables (for root biomass, litter decomposition, grass height, foliar C%,δ^15^C, δ^15^N) the direction of the net legume effect varied with laboratory.

**Fig. 2.**
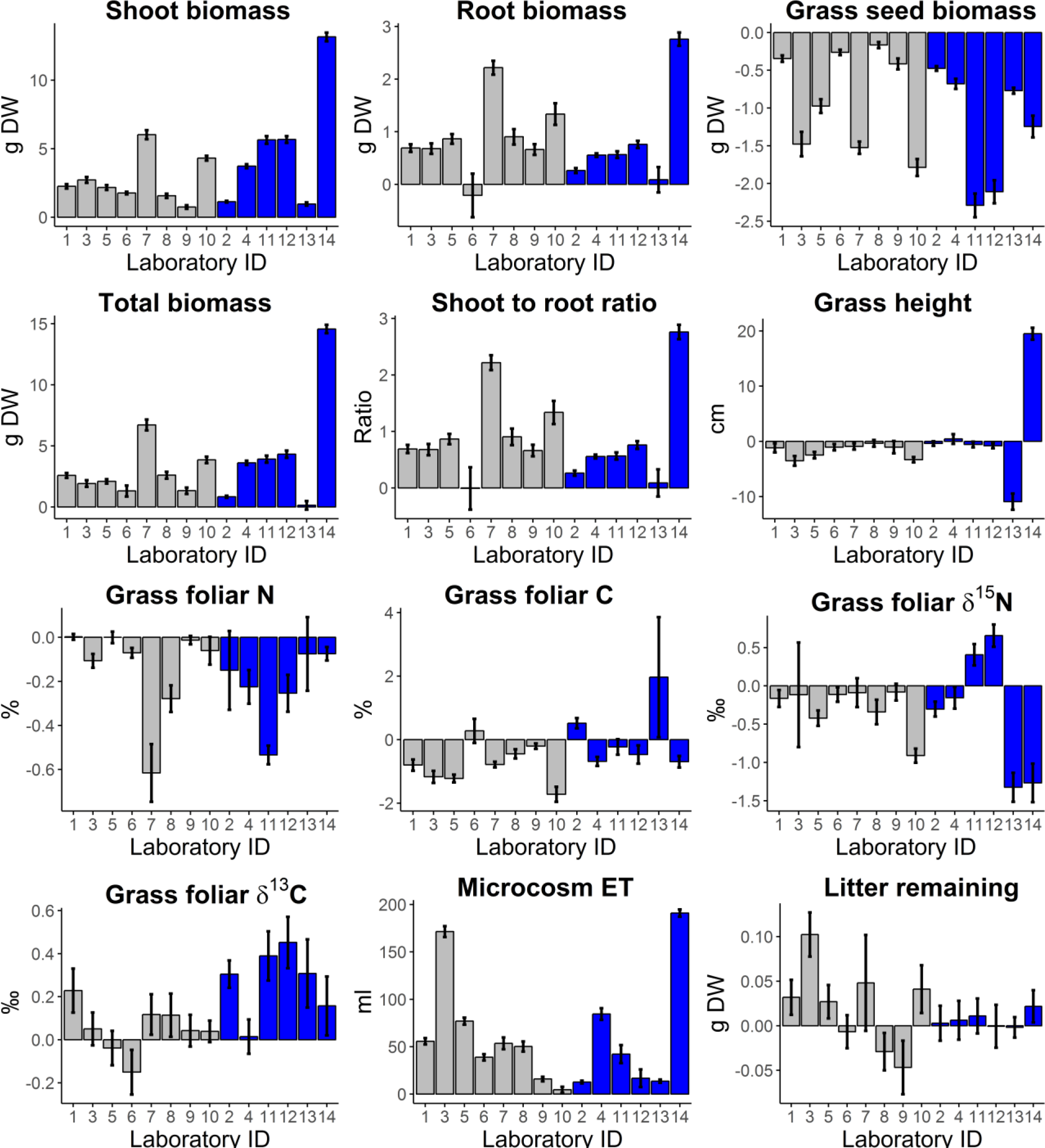
Net legume effect for the 12 response variables in 14 laboratories as affected by laboratory and SETUP (growth chamber vs. glasshouse) treatment. The grey and blue bars represent laboratories that used growth chamber and glasshouse set-ups, respectively. Bars show means by laboratory obtained by averaging over all CSV treatments, with error bars indicating ± 1 s.e.m. (n = 72 microcosms per laboratory).

Mixed-effects models testing the effect of the presence of legume species (LEG), laboratory (LAB), CSV, and their interactions (with experimental block—within-LAB growth chamber or glasshouse bench—as a random factor) on the 12 response variables revealed that for half of the variables the impact of the presence of legumes varied significantly with laboratory and CSV as indicated by the LEG×LAB×CSV three-way interaction (Table 1, Supplementary Figs 2 and 3). For the other half, significant two-way interactions between LEG×LAB and CSV×LAB were found. The same significant interactions were found for the analyses done on the first (PC1) and second (PC2) principal components from principal component PCA analysis that included all 12 response variables, which together explained 45% of the variation (Table 1; Supplementary Fig. 4ab). Taken together, these results suggest that the effect size and/or direction of the net legume effect was significantly different (i.e. not reproducible) in some laboratories and that the introduced CSV treatment affected reproducibility. In a complementary analysis including the SETUP in the model (and accounting for the LAB effect as a random factor), we found that the impact of the CSV treatment varied significantly with the SETUP (CSV×SETUP or LEG×CSV×SETUP interactions; Supplementary Table 3), suggesting the reproducibility of the results was different between glasshouses and growth chambers.

**Table 1.** Impact of experimental treatments on response variables. Mixed-effects model outputs summarizing the impacts of the presence of legumes (LEG), controlled systematic variability (CSV) and laboratory (LAB) on the 12 response variables. We also present the impact of experimental treatments on the first and second principal components (PC1 and PC2) of all 12 response variables. The response variables we measured are a typical ensemble of variables measured in plant-soil microcosm experiments (BM = biomass). † symbol indicates response variables measured for the grass *B. distachyon* only, whereas the rest of the variables were measured at the microcosm level, i.e. including the contribution of both the legume and the grass species. Stars indicate P-values (*** for *P* < 0.001; ** for *P* < 0.01; * for *P* < 0.05; + for *P* < 0.1; ns for *P* >0.1). DF = numerator degrees of freedom.

|  | DF | Shoot BM | Root BM | Seed BM<sup>†</sup> | Total BM | Shoot/Root | Grass height<sup>†</sup> | Shoot N%<sup>†</sup> |
| --- | --- | --- | --- | --- | --- | --- | --- | --- |
| LEG | 1 | 4602.95 (***) | 1131.64 (***) | 2186.64 (***) | 1022.28 (***) | 1137.01 (***) | 3.33 (+) | 366.73 (ns.) |
| CSV | 5 | 15.57 (***) | 23.93 (***) | 58.01 (***) | 2.70 (*) | 23.98 (***) | 23.36 (***) | 0.81 (ns.) |
| LAB | 13 | 1088.67 (***) | 182.53 (***) | 364.57 (***) | 1279.70 (***) | 183.42 (***) | 317.33 (***) | 350.91 (***) |
| LEG×CSV | 5 | 23.63 (***) | 4.48 (***) | 33.61 (***) | 5.69 (***) | 4.51 (***) | 2.62 (*) | 1.52 (ns) |
| LEG×LAB | 13 | 235.99 (***) | 40.58 (***) | 78.17 (***) | 126.70 (***) | 40.38 (***) | 49.89 (***) | 14.56 (***) |
| CSV×LAB | 65 | 6.54 (***) | 3.14 (***) | 6.93 (***) | 7.15 (***) | 3.16 (***) | 10.16 (***) | 1.97 (***) |
| LEG×LAB×CSV | 65 | 2.22 (***) | 1.12 (ns.) | 2.70 (***) | 1.14 (ns.) | 1.11 (ns.) | 1.45 (*) | 1.69 (***) |
|  |  | n = 1005 | n = 989 | n = 997 | n = 976 | n = 987 | n = 1008 | n = 939 |
| | DF | Shoot C%<sup>†</sup> | Shoot $\delta^{15}\text{N}^{\dagger}$ | Shoot $\delta^{13}\text{C}^{\dagger}$ | ET | Litter | PC1 | PC2 |
| LEG | 1 | 105.10 (***) | 14.42 (***) | 35.10 (***) | 1087.32 (***) | 1.81 (ns.) | 1229.11 (***) | 1005.88 (***) |
| CSV | 5 | 0.20 (ns.) | 8.84 (***) | 63.40 (***) | 20.13 (***) | 1.05 (ns.) | 12.86 (***) | 22.37 (***) |
| LAB | 1 | 175.31 (***) | 258.29 (***) | 577.80 (***) | 724.39 (***) | 117.34 (***) | 921.87 (***) | 516.00 (***) |
| LEG×CSV | 5 | 2.60 (*) | 6.48 (***) | 5.30 (***) | 4.73 (***) | 1.77 (ns.) | 46.99 (***) | 11.74 (***) |
| LEG×LAB | 1 | 11.90 (***) | 16.78 (***) | 2.30 (**) | 169.18 (***) | 2.05 (*) | 117.85 (***) | 28.02 (***) |
| CSV×LAB | 5 | 1.70 (**) | 4.39 (***) | 4.10 (***) | 20.54 (***) | 2.97 (***) | 7.20 (***) | 2.77 (***) |
| LEG×LAB×CSV | 5 | 1.32 (+) | 1.84 (***) | 1.01 (ns.) | 1.37 (*) | 1.17 (ns.) | 0.94 (ns.) | 1.65 (**) |
|  |  | n = 940 | n = 963 | n = 973 | n = 930 | n = 974 | n = 1008 | n = 1008 |

To answer the question of how many laboratories produced results that were statistically indistinguishable from one another (i.e. reproduced the same finding), we used Tukey’s post-hoc Honest Significant Difference (HSD) test for the LAB effect on the first and second principal components describing the net legume effect, which together explained 49% of the variation (Supplementary Fig. 4cd). Out of 14 laboratories, seven (PC1) or 11 (PC2) laboratories were statistically indistinguishable for controls; this value increased with either environmental or genotypic CSV for PC1 but not PC2 (Table 2). When we analyzed responses in growth chambers alone, five of eight laboratories were statistically indistinguishable for the control, but this increased to six out of eight in treatments with environmental CSV only and seven of eight in treatments with genotypic CSV (GEN_W_, GEN_A_ and GEN_W_+ENV_W_). In glasshouses, introducing CSV did not affect the number of statistically indistinguishable laboratories with respect to PC1 but decreased the number of statistically indistinguishable laboratories with respect to PC2 (Table 2).

**Table 2.**
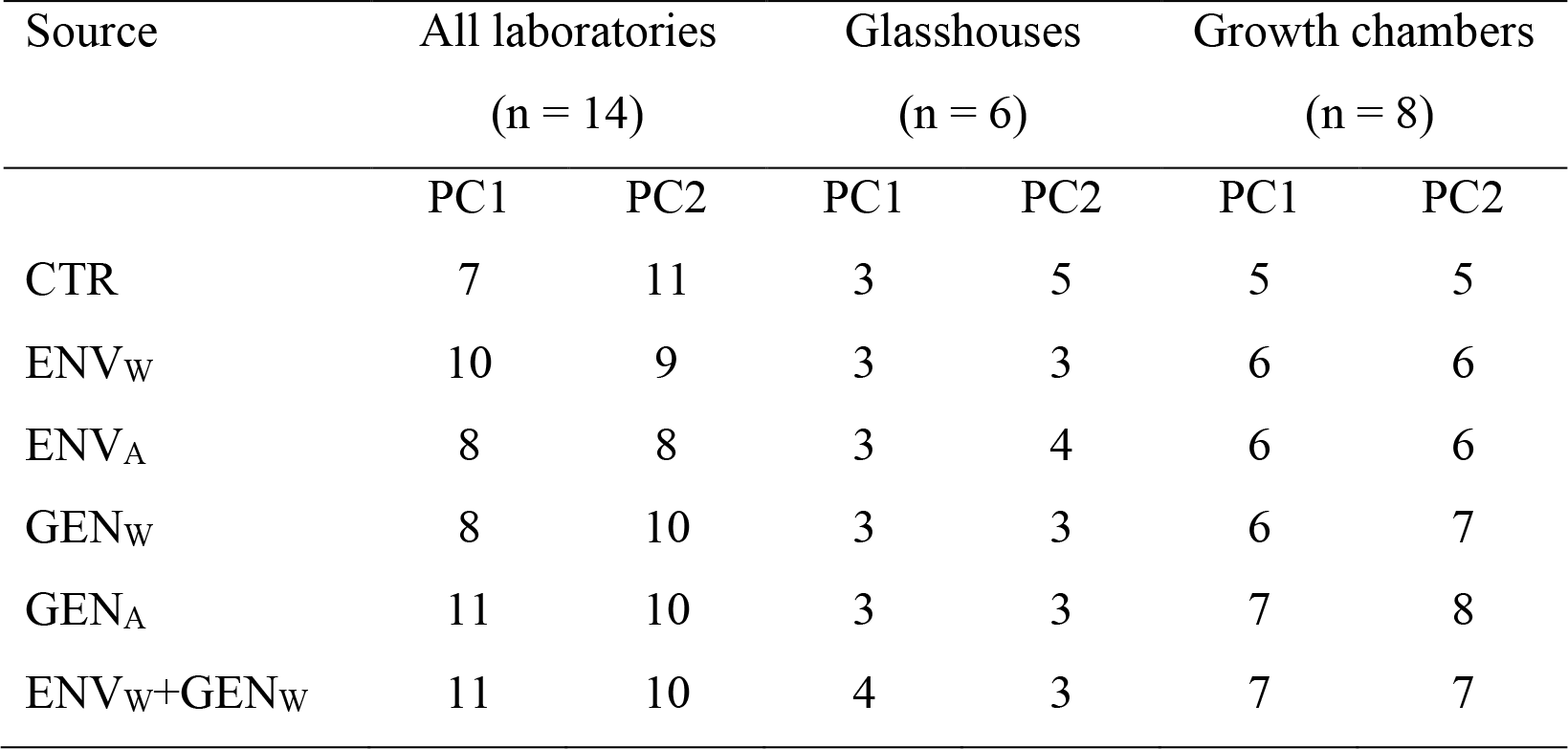
Impact of experimental treatments on the number of laboratories that reproduced the same finding. Numbers represent the total number of statistically indistinguishable laboratories based on a Tukey’s post-hoc Honest Significant Difference test of the first (PC1) and second (PC2) principal components of the net legume effect of the 12 response variables (see Supplementary Fig. 4cd for the PCA results). For a detailed description of experimental treatments and abbreviations see Fig. 1.

| Source | All laboratories<br>(n = 14) |  | Glasshouses<br>(n = 6) |  | Growth chambers<br>(n = 8) |  |
| --- | --- | --- | --- | --- | --- | --- |
|  | PC1 | PC2 | PC1 | PC2 | PC1 | PC2 |
| CTR | 7 | 11 | 3 | 5 | 5 | 5 |
| ENV<sub>W</sub> | 10 | 9 | 3 | 3 | 6 | 6 |
| ENV<sub>A</sub> | 8 | 8 | 3 | 4 | 6 | 6 |
| GEN<sub>W</sub> | 8 | 10 | 3 | 3 | 6 | 7 |
| GEN<sub>A</sub> | 11 | 10 | 3 | 3 | 7 | 8 |
| ENV<sub>W</sub>+GEN<sub>W</sub> | 11 | 10 | 4 | 3 | 7 | 7 |

We further assessed the impact of the experimental treatments on the among- and within-laboratory SD. Analysis of the among-laboratory SD of the net legume effect revealed a significant CSV×SETUP interaction (F_5,121_=7.38, P < 0.001) (Fig. 3a, b). This interaction included significantly lower fitted coefficients (i.e., lower among-laboratory SD) in growth chambers for GEN_W_ (*t*_5,121_ = −3.37, *P* = 0.001), GEN_A_ (*t*_5,121_ = −2.95, *P* = 0.004) and ENV_W_+GEN_W_ (*t*_1,121_ = −3.73, *P* < 0.001) treatments relative to CTR (see also Supplementary Note for full model output). For these three treatments, the among-laboratory SD of the net legume effect was 31.7% lower with genotypic CSV than without it, indicating increased reproducibility (Fig. 3a). The same analysis performed on within-laboratory SD of the net legume effect only found a slight but significant increase of within-laboratory SD in the GEN_A_ treatment (*t*_5,121_ = 3.52, *P* < 0.001) (Supplementary Note). We then tested whether there was a relationship between within- and among-laboratory SD with a statistical model for among-laboratory SD as a function of within-laboratory SD, SETUP, CSV and their interactions. We found a significant within-laboratory SD×SETUP×CSV three-way interaction (*F*_5,109_ = 2.4, *P* < 0.040) affecting among-laboratory SD (Supplementary Note). This interaction was the result of a more negative relationship between within- and among-laboratory SD in glasshouses relative to growth chambers, but with different slopes for the different CSV treatments (Fig. 4).

**Fig. 3.**
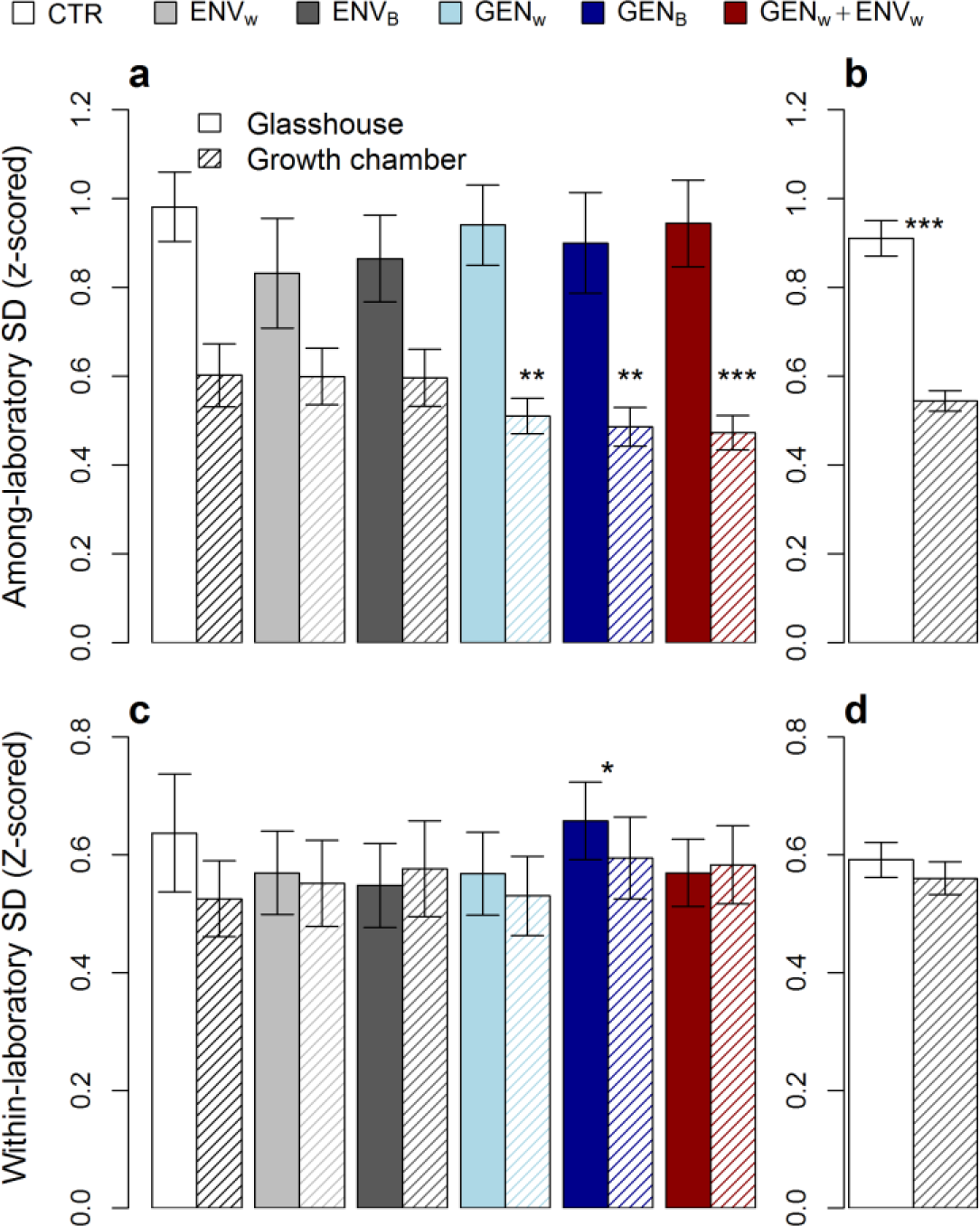
Among- and within-laboratory standard deviation (SD) of the net legume effect as affected by experimental treatments. Among-laboratory SD as affected by CSV and SETUP (**a**) and SETUP only (**b**). Within-laboratory SD as affected by CSV and SETUP (**c**) and SETUP only (**d**). Lower among-laboratory SD indicates enhanced reproducibility. Solid-filled bars and striped bars represent glasshouse (n = 6) and growth chamber setups (n = 8), respectively. Stars represent *P*-values (*** for *P* < 0.001, ** for *P* < 0.01, * for *P* < 0.05) indicating significantly different fitted coefficients according to the mixed-effects models (see Supplementary Notes for full model outputs); in (**c**) the star indicates the significant difference between GEN_A_ and CTR, irrespective of the type of SETUP. For a detailed description of experimental treatments and abbreviations see Fig. 1.

**Fig. 4.**
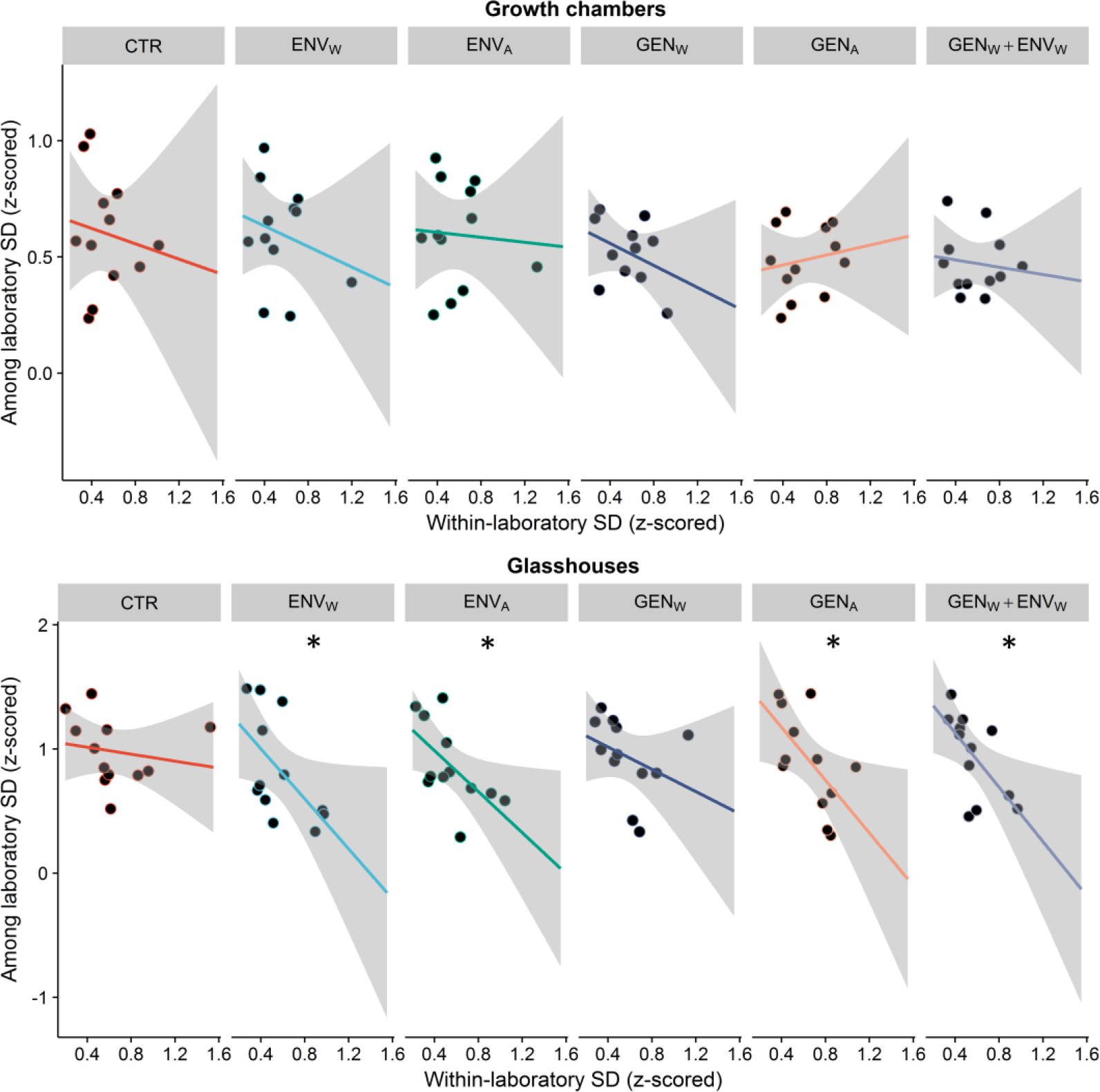
Relationship between within-laboratory SD and among-laboratory SD of the net legume effect as affected by experimental treatments. The figure illustrates the significant within-laboratory SD×SETUP×CSV three-way interaction (F_5,109_ = 2.4, P < 0.040) affecting among-laboratory SD (Supplementary Note). This interaction is the result of a more negative relationship between within- and among-laboratory SD in glasshouses relative to growth chambers, but with different slopes for the different CSV treatments. Points represent the 12 response variables. Stars represent *P* values < 0.05 for the individual linear regressions. Note the different scale for the y-axis between growth chambers and glasshouses. For a detailed description of experimental treatments and abbreviations see Fig. 1.

As we observed a tendency for CSV to increase within-laboratory variation (see Supplementary Note), we also analyzed the impact of the three CSV treatments that produced the most similar results (GEN_W_, GEN_A_, ENV_W_+GEN_W_) on the statistical power of detecting the net legume effect within individual laboratories. In growth chambers, adding genotypic CSV led to a slight reduction in statistical power relative to CTR (57% in CTR vs. 46% in the three treatments containing genotypic variability) that could have been compensated for by using eleven instead of six replicated microcosms per treatment. In glasshouses, owing to a higher effect size of the impact of the presence of legumes on the response variables, the statistical power for detecting the legume effect in CTR was slightly higher (68%) than in growth chambers, but was reduced to 51% on average for the three treatments containing genotypic CSV, a decrease that could have been compensated for by using 16 replicated microcosms instead of six.

Our findings provide compelling support for the hypothesis that introducing genotypic CSV in experimental designs can increase reproducibility of ecological studies^9–11^. However, the effectiveness of genotypic CSV for enhancing reproducibility varied with the setup as it only led to lower among-laboratory SD in growth chambers, not in glasshouses. Lower among-laboratory SD in growth chambers implies that the microcosms containing genotypic CSV were less strongly affected by unaccounted-for lab-specific environmental or biotic variables. Analyses performed at the level of individual variables (Table 1) showed that introducing genotypic CSV affected the among-laboratory SD in most but not all variables. This suggests that the relationship between genotypic CSV and reproducibility is probabilistic and results from the decreased likelihood that microcosms containing CSV will respond to unaccounted for lab-specific environmental factors in the same direction and with the same magnitude. The mechanism is likely to be analogous to the stabilizing biodiversity effect on ecosystem functions under changing environmental conditions^23–26^, but additional empirical evidence is needed to confirm this conjecture.

Introducing genotypic CSV increased reproducibility in growth chambers but not in glasshouses. Higher among-laboratory SD in glasshouses may indicate the existence therein of stronger laboratory-specific factors, and our deliberate use of different soils in the glasshouses presumably contributed to this effect. However, the among-laboratory SD in glasshouses decreased with increasing within-laboratory SD, irrespective of CSV, an effect that was less clear in growth chambers (Fig. 4). This observation is in line with the hypothesis put forward by Richter et al.^9^ that increasing the variance within experiments can reduce the among-laboratory variability of the mean effect sizes observed in each laboratory. However, the within-laboratory variability induced by the CSV treatments in glasshouses had no significant effect on among-laboratory variability, suggesting that our CSV treatments did not introduce sufficient within-microcosm variability to buffer against laboratory-specific factors for all response variables. This finding is in accordance with the two studies that explored the role of CSV for reproducibility in animal behavior and recommended the use of within-laboratory heterogenization to increase the likelihood of reproducibility of results across laboratories varying in experimental conditions^9,10^.

Our results also indicated that genotypic CSV was more effective in increasing reproducibility than environmental CSV, irrespective of whether the CSV was introduced within or among individual replicates (i.e., microcosms). However, we cannot discount the possibility that we found this result because our treatments with environmental CSV were less successful in increasing within-microcosm variability. Additional experiments could test whether other types of environmental CSV, such as soil nutrients, texture, or water availability, might be more effective at increasing reproducibility.

We expected higher overall productivity (i.e., a net legume effect) in the grass-legume mixtures and enhanced growth of *B. distachyon* because of the presence of the nitrogen (N)-fixing *M. truncatula*. However, these species were not selected because of their routine pairings in agronomic or ecological experiments (they are rarely used that way), but rather because they are frequently used in controlled environment experiments for functional genomics. Contrary to our expectation, and despite the generally lower ^15^N signature of *B. distachyon* in the presence of N-fixing *M. truncatula* (suggesting that some of the N fixed by *M. truncatula* was taken up by the grass), the biomass of *B. distachyon* was lower in the microcosms containing *M. truncatula*. Seed mass and shoot %N data of *B. distachyon* was lower in mixtures (Supplementary Fig. 1), suggesting that the two species competed for N. The lack of a significant N fertilization effect of *M. truncatula* on *B. distachyon* could have resulted from the asynchronous phenologies of the two species: the 8-10-week life cycle of *B. distachyon* may have been too short to benefit from the N fixation by *M. truncatula*. Whereas the direction of the legume effects were the same in the majority of laboratories, in 10% of the 168 laboratory × variable combinations (14 laboratories × 12 response variables) the direction differed from the among-laboratory consensus (Fig. 2).

Because well-established meta-analytical approaches can account for variation caused by local factors and still detect the general trends across different types of experimental setups, environments, and populations, we should ask whether the additional effort required for introducing CSV in experiments is worthwhile. Considering the current reproducibility crisis in many fields of science^27^, we suggest that it is, for at least three reasons. First, some studies become seminal without any attempts to reproduce them. Second, even if a seminal study that is flawed due to laboratory-specific biases is later proven wrong, it usually takes significant time and resources before its impact on the field abates. Third, the current rate of reproducibility is estimated to be as low as one-third^12–14^, implying that most data entering any meta-analysis are biased by unknown lab-specific factors. Addition of genotypic CSV may enhance the reproducibility of individual experiments and eliminate potential biases in data used in meta-analyses. Furthermore, if each individual study is less affected by laboratory-specific unknown environmental and biotic factors, then we would also need fewer studies to draw solid conclusions about the generality of phenomena. Therefore, we argue that investing more in making individual studies more reproducible and generalizable will be beneficial in both the short and long run. At the same time, adding CSV can reduce statistical power to detect experimental effects, so some additional experimental replicates would be needed when using it.

Overall, our study shows that results produced by microcosm experiments can be strongly affected by lab-specific factors. Although there are multiple causes for the reproducibility crisis^15,27,28^, deliberately including genetic variation in the studied organisms can be a simple solution for increasing the reproducibility of ecological studies performed in controlled environments. As the introduced genotypic variability only increased reproducibility in experimental setups with tightly controlled environmental conditions (i.e., in growth chambers using identical soil), future studies are needed to test whether the introduction of stronger sources of controlled within-laboratory variability can increase reproducibility in glasshouses with more loosely controlled environmental conditions and different soils.

## Acknowledgements

This study benefited from the CNRS human and technical resources allocated to the ECOTRONS Research Infrastructures and the state allocation ‘Investissement d’Avenir’ ANR-11-INBS-0001 as well as from financial support by the ExpeER (grant no. 262060) consortium funded under the EU-FP7 research program (FP2007-2013). *Brachypodium* seeds were kindly provided by Richard Sibout (Observatoire du Végétal, Institut Jean-Pierre Bourgin, F-78026 Versailles Cedex France) and *Medicago* seeds were supplied by Jean-Marie Prosperi (INRA Biological Resource Centre, F-34060 Montpellier Cedex 1, France). We further thank Jean Varale, Gesa Hoffmann, Paul Werthenbach, Oliver Ravel, Clement Piel and Damien Landais for assistance throughout the study. For additional acknowledgements see Supplementary Information.

## Author contributions

A.M. and J.R. designed the study with input from M.B, S.B and J-C.L. Substantial methodological contributions were provided by M.B., S.S., T.G., L.R. and M.S-L. Conceptual feedback on an early version was provided by G.F., N.E., J.R. and A.M.E. Data were analysed by A.M. with input from A.M.E. A.M. wrote the manuscript with input from all co-authors. All co-authors were involved in carrying out the experiments and/or analyses.

## Author Information

The authors declare no conflict of interest. Correspondence and request for materials should be addressed to A.M.

## METHODS

All laboratories tried to the best of their abilities to carry out an identical experimental protocol. Whereas not all laboratories managed to recreate precisely all details of the experimental protocol, we considered this to be a realistic scenario under which ecological experiments using microcosms are performed in glasshouses and growth chambers.

### Germination

The seeds from the three genotypes of *Brachypodium distachyon* (Bd21, Bd21-3 and Bd3-1) and *Medicago truncatula* (L000738, L000530 and L000174) were first sterilized by soaking 100 seeds in 100 mL of a sodium hypochlorite solution with 2.6% active chlorine, and stirred for 15 min using a magnet. Thereafter, the seeds were rinsed 3 times in 250 mL of sterile water for 10-20 seconds under shaking. Sterilized seeds were germinated in trays (10 cm deep) filled with vermiculite. The trays were kept at 4°C in the dark for three days before being moved to light conditions (300 μmol m^−2^ s^−1^ PAR) and 20/16°C and 60/70% air RH for day- and night-time, respectively. When the seedlings of both species reached 1 cm in height above the vermiculite, they were transplanted into the microcosms.

### Preparation of microcosms

All laboratories used identical containers (2-liter volume, 14.8-cm diameter, 17.4-cm height). Sand patches were created using custom-made identical “patch makers” consisting of six rigid PVC tubes (2.5 cm in diameter and 25 cm long), arranged in a circular pattern with an outer diameter of 10 cm. A textile mesh was placed at the bottom of the containers to prevent the spilling of soil through drainage holes. Filling of microcosms containing sand patches started with the insertion of the empty tubes into the containers. Thereafter, in growth chambers, 2000-g dry-weight of soil, subtracting the weight of the sand patches, was added into the containers and around the “patch maker” tubes. Because different soils were used in the glasshouses, the dry weight of the soil differed depending on the soil density and was first estimated individually in each laboratory as the amount of soil needed to fill the pots up to 2 cm from the top. After the soil was added to the containers, the tubes were filled with a mixture of 10% soil and 90% sand. When the microcosms did not contain sand patches, the amount of sand otherwise contained in the six patches was homogenized with the soil. During the filling of the microcosms, a common substrate for measuring litter decomposition was inserted at the center of the microcosm at 8 cm depth. For simplicity as well as for its fast decomposition rate, we used a single batch of commercially available tetrahedron-shaped synthetic tea bags (mesh size of 0.25 mm) containing 2 g of green tea (Lipton, Unilever), as proposed by the “tea bag index” method^20^. Once filled, the microcosms were watered until water could be seen pouring out of the pot. The seedlings were then manually transplanted to predetermined positions (Fig. 1), depending on the genotype and treatment. Each laboratory established two blocks of 36 microcosms each, resulting in a total of 72 microcosms per laboratory, with blocks representing two distinct chambers in growth chamber setups or two distinct growth benches in the same glasshouse.

### Soils

All laboratories using growth chamber setups used the same soil, whereas the laboratories using glasshouses used different soils (see Supplementary Table 1 for the physicochemical properties of the soils). The soil used in growth chambers was classified as a nutrient-poor cambisol and was collected from the top layer (0–20 cm) of a natural meadow at the Centre de Recherche en Ecologie Expérimentale et Prédictive—CEREEP (Saint-Pierre-Lès-Nemours, France). Soils used in glasshouses originated from different locations. The soil used by laboratory L2 was a fluvisol collected from the top layer (0-40 cm) of a quarry site near Avignon, in the Rhône valley, Southern France. The soil used by laboratory L4 was collected from near the La Cage field experimental system (Versailles, France) and was classified as a luvisol. The soil used by labs L11 and L12 was collected from the top layer (0-20cm) within the haugh of the river Dreisam in the East of Freiburg, Germany. This soil was classified as an umbric gleysol with high organic carbon content. The soil from laboratory L14 was classified as a eutric fluvisol and was collected on the field site of the Jena Experiment, Germany. Prior to the establishment of microcosms, all soils were air-dried at room temperature for several weeks and sieved with a 2-mm mesh sieve. A common inoculum was provided to all laboratories to assure that rhizobia specific to *M. truncatula* were present in all soils.

### Abiotic environmental conditions

The set points for environmental conditions were 16 h light (at 300 μmol m^−2^ s^−1^ PAR) and 8h dark, 20/16°C, 60/70% air RH for day- and night-time, respectively. Different soils (for glasshouses) and treatments with sand patches likely affected water drainage and evapotranspiration. The watering protocol was thus based on drying weight relative to weight at full water holding capacity (WHC). The WHC was estimated based on the weight difference between the dry weight of the containers and the wet weight of the containers 24 h after abundant watering (until water was flowing out of the drainage holes in the bottom of each container). Soil moisture was maintained between 60 and 80% of WHC (i.e. the containers were watered when the soil water dropped below 60% of WHC and water added to reach 80% of WHC) during the first 3 weeks after seedling transplantation and between 50 and 70% of WHC for the rest of the experiment. Microcosms were watered twice a week with estimated WHC values from two microcosms per treatment. To ensure that the patch/heterogeneity treatments did not become a water availability treatment, all containers were weighed and brought to 70 or 80% of WHC every two weeks. This operation was synchronized with within-block randomization. All 14 experiments were performed between October 2014 and March 2015.

### Sampling and analytical procedures

After 80 days, all plants were harvested. Plant shoots were cut at the soil surface, separated by species, and dried at 60ºC for three days. Roots and any remaining litter in the tea bags were washed out of the soil using a 1-mm mesh sieve and dried at 60ºC for three days. Microcosm evapotranspiration rate was measured before the harvesting as the difference in weight changes from 70% of WHC after 48 h. Shoot C%, N%, δ^13^C, and δ^15^N were measured on pooled shoot biomass (including seeds) of *B. distachyon* and analyzed at the Göttingen Centre for Isotope Research and Analysis using a coupled system consisting of an elemental analyzer (NA 1500, Carlo Erba, Milan, Italy) and a gas isotope mass spectrometer (MAT 251, Finnigan, Thermo Electron Corporation, Waltham, Massachusetts, USA).

### Data analysis and statistics

All analyses were done using R version 3.2.4^29^. Data from each laboratory were first screened individually for outliers, and values that were lower or higher than three times the inter-quartile range (representing less than 1.7% of the whole dataset) were removed and considered missing values. We then assessed whether the impact of the presence of legume (LEG) varied with laboratory (LAB) and the treatment of controlled systematic variability (CSV). This was tested individually for each response variable (Table 1) with a mixed-effects model using the “nlme” package^30^. Following the guidelines suggested by Zuur et al. (2009)^31^, we first identified the most appropriate random structure using a restricted maximum likelihood (REML) approach and selected the random structure with the lowest Akaike information criterion (AIC). For this model, CSV and LAB were included as fix factors, experimental block as a random factor, and a “varIdent” weighting function to correct for heteroscedasticity resulting from more heteroscedastic data at the LAB and LEG level (R syntax: “model= lme (response variable ~ LEG*CSV*LAB, random=~1|block, weights=varIdent (form = ~1|LAB*LEG)”) (Table 2). As the LAB and SETUP experimental factors were not fully crossed (i.e. laboratories performed the experiment only in one type of setup), the two experimental variables could not be included simultaneously as fixed effects. Therefore, to test for the SETUP effect, we used an additional complementary model including CSV and SETUP as fix effects and laboratory as a random factor (R syntax: “model= lme (response variable ~ LEG*CSV*SETUP, random=~1|LAB/block, weights=varIdent (form = ~1|LAB*LEG)”) (Supplementary Table 3). To test whether the results were affected by the collinearity among the response variables, the two models also were run on the first (PC1) and second (PC2) principal components the 12 response variables (Fig. 4ab). PCs were estimated using the “FactoMineR” package^32^, with missing values replaced using a regularized iterative multiple correspondence analysis^33^ in the “missMDA” package^34^. The same methodology was used to compute a second PCA derived from the net legume effect on the 12 response variables (Supplementary Fig. 4cd). To assess how many laboratories produced results that were statistically indistinguishable from one another, we applied Tukey’s post-hoc HSD test in the “multcomp” package to lab-specific estimates of PC1 and PC2 (Table 2).

To assess how the CSV treatments affected the among- and within-laboratory variability, we used the standard deviation (SD) instead of the coefficient of variation, because the net legume effect contained both positive and negative values. To calculate among- and within-laboratory SDs, we centered and scaled the raw values using the z-score normalization [*z*-scored variable = raw value–mean)/SD] individually for each of the 12 response variables. Among-laboratory SD was computed from the mean of the laboratory *z*-scores for each response variable, CSV, and SETUP treatments (n = 144; 6 CSV levels × 2 SETUP levels × 12 response variables). Within-laboratory SDs were computed from the values measured in the six replicated microcosms for each CSV and SETUP treatment combination, individually for each response variable, resulting in a dataset with the same structure as for among-laboratory SDs (n = 144; 6 CSV levels × 2 SETUP levels × 12 response variables). Some of the 12 response variables were intrinsically correlated, but most had correlation coefficients < 0.5 (Supplementary Fig. 5) and were therefore treated as independent variables. To analyze and visualize the relationships between the SDs calculated from variables with different units, before the calculation of the among- and within-laboratory SD, the raw values of the 12 response variables were centered and scaled.

The impact of experimental treatments on among- and within-laboratory SD was analyzed using mixed-effect models, following the same procedure described for the individual response variables. The model with the lowest AIC included a random slope for the SETUP within each response variable as well as a “varIdent” weighting function to correct for heteroscedasticity at the variable level (R syntax: “model= lme (SD ~ CSV*SETUP, random=~SETUP|variable, weights=varIdent (form = ~1|variable)) (see also Supplementary Notes). The relationship between within- and among-laboratory SD also was tested with a model with similar random structure but with among-laboratory SD as a dependent variable and within-laboratory SD, CSV and SETUP as predictors.

Because the treatments containing genotypic CSV increased reproducibility in growth chambers, but slightly increased within-laboratory SD, we also examined the effect of adding CSV on the statistical power for detecting the net legume effect in each individual laboratory. This analysis was done with the “power.anova.test*”* function in the “base” package. We computed the statistical power of detecting a significant net legume effect (if one had used a one-way ANOVA for the legume treatment) for CTR, GEN_W_, GEN_A_ and ENV_W_+GEN_W_ treatments for each laboratory and response variable. This allowed us to calculate the average statistical power for the aforementioned treatments and how many additional replicates would have been needed to achieve the same statistical power as we had in the CTR.

The data that support the findings of this study are available from the corresponding author upon reasonable request.

